# Cholinergic dynamics in the septo-hippocampal system provide phasic multiplexed signals for spatial novelty and correlate with behavioral states

**DOI:** 10.1101/2025.01.21.634097

**Authors:** Fatemeh Farokhi Moghadam, Blanca E. Gutierrez Guzman, Xihui Zheng, Mina Parsa, Lojy M. Hozyen, Holger Dannenberg

**Affiliations:** Department of Bioengineering, George Mason University, Fairfax, VA, United States; Interdisciplinary Program for Neuroscience, George Mason University, Fairfax, VA, United States

**Keywords:** medial septum, hippocampal formation, acetylcholine, encoding and retrieval, novel object location, fiber photometry

## Abstract

In the hippocampal formation, cholinergic modulation from the medial septum/diagonal band of Broca (MSDB) is known to correlate with the speed of an animal’s movements at sub-second timescales and also supports spatial memory formation. Yet, the extent to which sub-second cholinergic dynamics, if at all, align with transient behavioral and cognitive states supporting the encoding of novel spatial information remains unknown. In this study, we used fiber photometry to record the temporal dynamics in the population activity of septo-hippocampal cholinergic neurons at sub-second resolution during a hippocampus-dependent object location memory task using ChAT-Cre mice. Using a general linear model, we quantified the extent to which cholinergic dynamics were explained by changes in movement speed, behavioral states such as locomotion, grooming, and rearing, and hippocampus-dependent cognitive states such as recognizing a novel location of a familiar object. The data show that cholinergic dynamics contain a multiplexed code of fast and slow signals i) coding for the logarithm of movement speed at sub-second timescales, ii) providing a phasic spatial novelty signal during the brief periods of exploring a novel object location, and iii) coding for environmental novelty at a seconds-long timescale. Furthermore, behavioral event-related phasic cholinergic activity around the onset and offset of the behavior demonstrates that fast cholinergic transients help facilitate a switch in cognitive and behavioral state before and during the onset of behavior. These findings enhance understanding of the mechanisms by which cholinergic modulation contributes to the coding of movement speed and encoding of novel spatial information.

## INTRODUCTION

Cognitive map-based navigation requires the encoding, storage, and recall of spatial information to navigate environments effectively (Schiller et al., 2015). This process integrates sensory inputs, motor planning, and memory, relying on a mental representation of both current and future locations (Hinman et al., 2018). The hippocampal formation plays a central role in spatial navigation by supporting the formation and retrieval of spatial memories (Eichenbaum, 2017). Cholinergic projections to the hippocampal formation arise from cholinergic neurons in the medial septum and diagonal band of Broca (MSDB) (Dutar et al., 1995). Changes in cholinergic modulation by medial septal projection neurons can cause changes in brain and behavioral states (Alonso and Köhler, 1984; Dannenberg et al., 2017; Hinman et al., 2018; Mesulam et al., 1983; Rye et al., 1984). In the hippocampus, optogenetic stimulation of cholinergic projection neurons induces theta and gamma power in anesthetized mice (Dannenberg et al., 2015) and enhances theta rhythmic activity while suppressing sharp wave-ripples (SPW-Rs) in freely behaving mice (Vandecasteele et al., 2014). These findings support the influential hypothesis that acetylcholine (ACh) enhances processing of feedforward sensory input to the cortex to promote the encoding of novel memories while suppressing retrieval of previously stored memories via feedback excitation (Hasselmo, 2006; Hasselmo et al., 1995). Conversely, during quiet wakefulness and non-REM sleep, cholinergic tone is reduced, allowing feedback excitation to dominate during SPW-Rs serving memory consolidation and action planning (Barry et al., 2012; Fernández-Ruiz et al., 2019; Hasselmo, 2006; Hasselmo and McGaughy, 2004; Rogers and Kesner, 2003; Zhang et al., 2024, 2021).

While neuromodulation by ACh in the hippocampal formation has been linked to the encoding of spatial memories by modulating network dynamics, synaptic plasticity, and neuronal excitability (Blokland et al., 1992; Dannenberg et al., 2017; Hasselmo, 2006; Hasselmo et al., 2017; Hasselmo and McGaughy, 2004; Ohno et al., 1993; Stancampiano et al., 1999; Záborszky et al., 2018), recent mouse studies suggest that cholinergic signals may provide a code for movement speed and promote processing of sensory information associated with exploratory behavior, as opposed to modulating memory function (Cassity et al., 2023; Dannenberg et al., 2015; Kopsick et al., 2022). In addition, cholinergic modulation has been suggested to provide a novelty signal to distinguish between familiar and novel spatial information (Gómez-Ocádiz et al., 2022). Yet, how cholinergic dynamics in the septo-hippocampal system can support those multiple cognitive and sensory-motor functions is largely unknown. Furthermore, whether cholinergic dynamics are fast enough to provide novelty signals that can support the encoding of novel spatial information during active exploratory behavior remains largely unknown. While cholinergic signals can act across different temporal scales (Disney and Higley, 2020; Sarter and Lustig, 2020), the timescales of cholinergic dynamics in mice performing a hippocampus-dependent memory task have not been quantified.

Traditionally, cholinergic signaling was thought to be primarily slow and diffuse, with ACh released in a tonic manner across large brain regions to regulate broad functions like arousal. This view emerged largely from techniques such as microdialysis, which lack the temporal resolution and measures changes over minutes (Jiménez-Capdeville and Dykes, 1996; Kametani and Kawamura, 1991; Marrosu et al., 1995). Recent research, however, has shown that phasic and tonic ACh signaling operate on both fast (sub-seconds to seconds) and slow (minutes to hours) timescales (Disney and Higley, 2020). In cortical regions, tonic Ach release is associated with changes in brain and behavioral states over slow timescales, while phasic Ach release integrates sensory and motor information related to behavioral events at fast timescales (Cassity et al., 2023; Dannenberg et al., 2017, 2016; Disney and Higley, 2020; Dudar et al., 1979; Gritton et al., 2016; Sarter and Lustig, 2020; Yogesh and Keller, 2024; Záborszky et al., 2018; Zhu et al., 2023)

Behavioral studies have shown that manipulations inducing or inhibiting rapid cholinergic transients directly affect task performance, including attention and cue detection (Gritton et al., 2016; Howe et al., 2017, 2010). While a multiplexed combination of fast and slow cholinergic signals has been demonstrated in the auditory cortex (Kuchibhotla et al., 2017), it remains unclear whether cholinergic modulation in the septo-hippocampal system can provide a multiplexed combination of fast and slow signals, potentially supporting memory and motor-sensory tasks.

In this study, we used fiber photometry to record the population activity of cholinergic neurons in the MSDB at sub-second timescales. These recordings were performed using Chat-Cre mice engaged in an object location memory task (ObLoM). Mice were video tracked to quantify movement speed and behaviors. The results show that cholinergic activity in the septo-hippocampal circuitry exhibits multiplexed coding: fast cholinergic dynamics provide a code for movement speed and transient spatial novelty signals during the brief moments of object exploration at novel but not familiar locations. In contrast, slow changes in cholinergic activity are associated with the novelty of environmental context. These data contribute to our understanding of the mechanisms by which cholinergic modulation contributes to the encoding of spatial memories, including novel object-space associations.

## MATERIALS AND METHODS

### Animals

We used a total of six transgenic heterozygous mice (see table 1) expressing Cre-recombinase under the control of the choline-acetyltransferase promoter (ChAT-IRES-Cre) at the age of 3-8 months. Breeder pairs were purchased from the Jackson Laboratory (C57BL/6J X B6;129S6Chat^TM2(cre)Lowl^/J, 3 mice; C57BL/6J X B6;129S6Chat^TM1(cre)Lowl^/J, 3 mice). This study includes data from three mice which were also used to collect additional data for a previously published study (Kopsick et al., 2022). Prior to surgery, mice were housed in Plexiglas cages with their siblings. Post-surgery, they were individually housed in extended-height cages on a 12-hour reversed light/dark cycle with food and water available ad libitum. The cages included a spherical treadmill for enrichment, physical exercise, and stress relief. All mice were handled and habituated to the experimenter and testing room prior to the start of experiments. Experimental procedures for newly acquired data adhered to the regulations outlined in the Guide for the Care and Use of Laboratory Animals by the National Research Council and approved by the Institutional Animal Care and Use Committee of George Mason University (Protocol No.: 0501, approved on January 24, 2022).

**TABLE 1.** Information on mice used in experiments. GMU, George mason university; BU, Boston university; ObLoM, Object location memory task.

| Mouse ID | Lab | Gender | Age (month) | Number of trials | Genotype | GCaMP | Performance of ObLoM task (days after surgery) |
| --- | --- | --- | --- | --- | --- | --- | --- |
| #1 | GMU | Male | 8 | 1 | ChAT-Cre(tg/+) | jGCaMP 8m | 12 |
| #2 | GMU | Male | 6 | 2 | ChAT-Cre(tg/+) | jGCaMP 8s | 15, 21 |
| #3 | GMU | Male | 4 | 1 | ChAT-Cre(tg/+) | jGCaMP 8s | 12 |
| #611915 | BU | Female | 3–6 | 2 | ChAT-Cre(tg/+) | jGCaMP 7s | 10, 14 |
| #611916 | BU | Female | 3–6 | 2 | ChAT-Cre(tg/+) | jGCaMP 7s | 11, 16 |
| #611926 | BU | Male | 3–6 | 2 | ChAT-Cre(tg/+) | jGCaMP 7s | 11, 15 |

### Surgery

Anesthesia was induced with 4% isoflurane in a gas chamber and maintained at 1-2% (mixed with 100% oxygen) via the nose cone on the stereotaxic instrument. The body temperature was kept steady and monitored throughout the surgery using a heating pad and homeothermic monitoring system (36-37°C). Preoperative care included administering atropine (0.1 mg/kg, s.c.), ketoprofen (5 mg/kg, s.c.), enrofloxacin (7.5 mg/kg, s.c.), and local anesthetic (lidocaine 0.5%, 5 mg/kg) underneath the scalp before making the surgical incision.

#### Virus injection

Craniotomy was performed for a recombinant adeno-associated virus (rAAV) injection of calcium indicators from the GCaMP family (jGCaMP7s, jGCaMP8s, or jGCaMP8m; see Table 1) and for the chronic implantation of an optical fiber above the MSDB. Both procedures use the same coordinates from on the Paxinos and Franklin atlas (Paxinos and Franklin, 2019): 1.05 mm anterior from Bregma and 0.7 mm lateral to Bregma. The injection needle (Table 2) was lowered into the MSDB at an 8° polar angle and a –90° azimuth angle, and total volume of 1000 nl of virus solution was injected at a rate of 100 nl/min at two ventral sites, –4.8 mm and –4.4 mm below the skull surface (500 nl at each), using an injection pump (Table 2). The needle for virus injection was left in place for 10 minutes after each injection to ensure diffusion and prevent backflow of the virus solution.

**TABLE 2.** Materials and resources.

| Material | Specifications |
| --- | --- |
| Injection needle | NanoFil Needle NF34BV ,34G, beveled, WPI |
| Injection pump | UMP3T-1; MICRO2T SMARTouch™ controller, WPI |
| Implantable optical fiber | MFC_200/250-0.66_7mm_ZF1.25(G)_FLT; Doric lenses |
| Fiber Photometry System | RZ10X, Tucker Davis Technologist |
| Mono fiber optic patch cord | MFP_200/230/900-0.57_2.5m_FC-MF1.25(F)_LAF, Doric Lenses |
| Rotary joint | FRJ_1x1_FC-FC, Doric lenses |
| Dulbecco's phosphate-buffered saline (DPBS) | Catalog#14040133, ThermoFisher Scientific |
| Vibratome | VT1000 S, Leica |
| Goat anti-ChAT affinity purified polyclonal antibody | Catalog #AB 144P, Merck Millipore |
| Triton X-100 detergent | X100-100ML, Sigma-Aldrich, St. Louis, MO |
| Cy3-conjugated donkey anti-goat IgG polyclonal antibody | Catalog #AP180C, Merck Millipore |
| Aqua-Poly/Mount | #18606-20, Polysciences, Inc., Warrington, PA |
| Confocal laser scanning microscope | LSM 800, Zeiss |

#### Implantation of optical fiber

An optical fiber (total diameter: 250 µm; 200 µm core; N.A. 0.66, see Table 2) was implanted at the same coordinates and angle used for the virus injection. The fiber was lowered 4.2 mm from the skull surface and secured to the animal’s skull with black dental cement. Four stainless steel anchoring screws were implanted into the skull to reinforce the attachment of the dental cement and optical fiber. Care was taken that the anchoring screws did not protrude into brain tissue.

#### Post-Surgical Care

Mice received ketoprofen (5 mg/kg, s.c.) and enrofloxacin (7.5 mg/kg, s.c.) for two days after surgery. They were given one week for full recovery before the start of experiments.

### Fiber photometry

Newly acquired data were collected with TDT’s fiber photometry system (Tucker Davis Technologist; RZ10X), which is similar to the previous custom-made system used in (Kopsick et al., 2022) but provided several improvements with respect to the signal-to-noise ratio, including a more stable non-magnetic connection to the implanted optical fiber to reduce movement noise. The TDT system used lock-in amplification, configured with a 465 nm LED driver modulated at 211 Hz for Ca^2+^-dependent excitation of GCaMP and a 405 nm UV LED driver modulated at 531 Hz for excitation of GCaMP at its isosbestic point. Laser light was transmitted into the brain using fiber optic patch cords connected through a rotary joint (Table 2). The power of the laser entering the implanted optical fiber was assessed both before and after each recording session and adjusted to deliver approximately 40 µW of laser power into the MSDB. The data were demodulated with a 6^th^-order low-pass filter set to 6 Hz. LED currents were adjusted to return a voltage between 5 and 10 mV for each signal, offset by 5 mA. Data were acquired at a sampling rate of 610 Hz. TDT Synapse software was used for data acquisition.

### Behavioral tests and video-tracking

Data were acquired from animals performing an ObLoM, which relies on spontaneous behavior. The behavioral tests were performed in a dimly lit room dedicated for behavioral experiments with no windows to minimize external stimuli.

#### Arena and objects

The recording arena consisted of an open field box made of black acrylic, measuring 40 x 40 cm² with 30 cm high walls. A visual cue card (a white triangle) was placed on one wall, the same wall across all experiments. Two identical cylindrical objects (50 ml Conical Tubes) were placed 10-cm away from two adjacent or non-adjacent corners of the arena in the Sample and Test phases, respectively.

#### Object location memory task

The object location memory (ObLoM) task was adapted from (Schlesiger et al., 2021). We conducted one or two trials of the ObLoM task in each of the 6 mice, each comprising a Sample and Test phase. Sample and Test phases were 15-min long and were separated by a 1-hour delay period, during which the mice were returned to their home cage. To prevent olfactory interference and preference for one object, the maze and objects were cleaned with 70% isopropanol between phases. In four out of six mice (see Table 1), sessions were repeated in a counterbalanced design with the stationary and non-stationary objects reversed to control for potential location preferences of mice. The interval between recording sessions for each mouse ranged from 4 to 6 days (Mean ± Std = 4.75 ± 0.95). Before the start of the ObLoM task, mice were habituated to the experimenter, the recording room, and the maze, during which the mice were allowed to explore the open field box for at least 20 minutes daily for 5 days.

#### Data acquisition

Fiber photometry data were acquired in continuous recordings during all phases of the memory task (Sample phase, delay period, and Test phase), within a time window of 5 days to 3 weeks after the virus injection.

#### Video-tracking

Mice were video-tracked using a camera ceiling-mounted above the arena. The sampling rate ranged from 10-30 frames per second. All recordings were subsequently resampled via Fourier-based interpolation to 30 Hz for consistent comparison. The camera and the fiber photometry system were synchronized using TTL pulses.

### Histology

#### Tissue preparation

Mice were deeply anesthetized with isoflurane and transcardially perfused with Dulbecco’s phosphate-buffered saline (DPBS) (Table 2) containing 0.9% calcium chloride, followed by 10% buffered formalin. Mice were decapitated post perfusion, and the heads were stored in formalin for 24-48 hours at 4 °C for further tissue fixing. Brains were then extracted and stored in DPBS at 4°C.

#### Sectioning and Staining

30 µm coronal slices of the medial septum were collected in DPBS using a vibratome (Table 2). Immunohistochemical staining for choline acetyltransferase (ChAT) in the MSDB was performed as described previously (Kopsick et al., 2022). Slices were washed three times for 15 minutes with DPBS containing 0.9% calcium chloride, then incubated with a goat anti-ChAT affinity purified polyclonal antibody (Table 2), diluted 1:200 with 0.3% Triton X-100-DPBS (Table 2) for 2 days at 4°C. For the secondary antibody incubation, after washing three times with DPBS, slices were incubated for 2 hours at room temperature with a Cy3-conjugated donkey anti-goat IgG polyclonal antibody (Table 2), diluted 1:200. Finally, slices were washed three more times, mounted on glass slides with Aqua-Poly/Mount (Table 2).

#### Imaging

Slides were imaged using a confocal fluorescent microscope (Table 2) with a suitable objective (10X and 20X). Optical fiber placements were histologically verified by visualizing fiber tracks using the mouse atlas (Paxinos and Franklin, 2019) and checking the alignment of GCaMP expression with regions containing ChAT-positive neurons.

### Analysis

#### Fiber photometry signal

Signal processing was performed as reported previously(Kopsick et al., 2022). Briefly, the recorded calcium and isosbestic control signals were resampled to match the video frame rate (30 Hz). We accounted for photobleaching and motion artifacts in the fiber-photometry signal by subtracting an adjusted control signal from the calcium signal. To compute the adjusted control signal, we first subtracted the isosbestic control signal from the calcium signal, fitted a second-degree curve to the result of that subtraction, and added the fitted curve back to the isosbestic control signal. In a second step, we adjusted the adjusted control signal’s amplitude so that the correlation with the calcium signal was maximal. Specifically, we created an optimization task to determine the values of two parameters, *⍺* and *β*, minimizing the expression Σ(s − (c ∗ α + β))^2^, where *s* represents the main signal, and *c* represents the adjusted control signal. Lastly, we computed 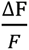 as 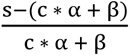.

#### Calculation of movement speed using markerless pose estimation

We utilized DeepLabCut (Mathis et al., 2018), a deep learning tool, for markerless pose estimation of mice’s body parts, including the neck and tail base. To estimate the animal’s movement speed, we tracked the neck position in the maze.

#### Speed tuning across different time scales

To determine the time scale at which changes in movement speed and cholinergic activity aligned best, we calculated the Pearson correlation coefficient between the logarithm of the animal’s movement speed and the cholinergic signal measured as 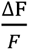 (Dannenberg et al., 2019) using logarithmically increasing smoothing window sizes ranging from 0.25 s to 256 s.

#### Classification of behavioral profiles

We utilized DeepEthogram (Bohnslav et al., 2021) to classify behavioral profiles in mice during experimental sessions. For our study, we selected a random 2– 3-minute window from each video and manually labeled the frames based on a predefined list of behaviors to create the training dataset. We then trained DeepEthogram’s algorithm to label the remaining video portions. The output provided a list of frame numbers with binary indicators (1 or 0) signifying the presence or absence of each behavior. We focused on classifying behaviors including locomotion, rearing, grooming, and object exploration. Locomotion refers to any movement in which the mouse changes its body position in space. Rearing was defined as any instance of the mice standing on their hind legs, either supported or unsupported by a wall or an object. Grooming includes any self-maintenance behavior, such as licking fur, scratching, or face washing. Object exploration included any interaction with an object, such as sniffing the object, moving around the object, stretched attend-behavior toward the object, or rearing supported by the object. To differentiate between exploring stationary and non-stationary objects in the Test phase, we used data extracted from DeepLabCut. If the animal’s nose was within a 10 cm radius around the center of the stationary or non-stationary object, we classified the timepoints as exploring the stationary and non-stationary object, respectively. Finally, the experimenter confirmed all labels and corrected errors made by DeepLabCut and DeepEthogram to improve accuracy of the exact start and end times of each behavior.

#### Discrimination index

To quantify the animal’s performance in the object location memory task, i.e., the ability to distinguish between a familiar and novel spatial location of objects, we used the discrimination index (DI). The DI index was calculated as

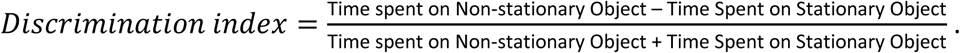

#### General linear model and normalization

To differentiate between the individual effects of each behavior on cholinergic activity, we employed a general linear model (GLM):

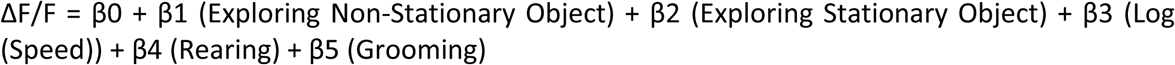

ΔF/F = β0 + β1 (Exploring Non-Stationary Object) + β2 (Exploring Stationary Object) + β3 (Log (Speed)) + β4 (Rearing) + β5 (Grooming)

However, the signal-to-noise ratio in the fiber photometry signal differs across discontinuous sessions. To analyze discontinuous fiber photometry signals across sessions, we leveraged the consistent correlation of cholinergic activity to the animal’s movement speed and expressed each model coefficient as multiples of the coefficient for movement speed.

#### Event-triggered cholinergic activity

For each behavior, we extracted the start and end times and analyzed the z-score of cholinergic activity measured as 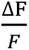 during the start and end times as well as in the 5-s before and the 5-s after the start and end of the behavior, respectively. We removed any short bouts (<2 s) and excluded bouts if the same behavior occurred within 4 s before the onset or after the offset of the bout. To average event-triggered cholinergic activity across events, we standardized the data points between start and end by linearly time-warping the cholinergic activity to a uniform length. This approach allowed us to compute an event-triggered temporal profile of cholinergic dynamics for each behavior.

### Experimental Design and Statistical Analysis

Data analysis and statistical tests were conducted using custom-written Python scripts and DATAtab (https://datatab.net), with all codes and processed data accessible via the GitHub repository (https://github.com/dannenberglab/Cholinergic-dynamics-in-spatial-exploration). Cholinergic activity and movement speed were smoothed using a 0.5-second moving average, and predicted cholinergic activity based on movement speed was calculated using a linear regression model. Correlations between time-series data were assessed using Pearson’s correlation coefficient. A GLM was applied using built-in Python functions to evaluate the effects of behavioral signals and movement speed on cholinergic activity, while a Linear Mixed Model (LMM) analyzed differences in correlations and the effects of object exploration between Sample and Test sessions. Observed cholinergic activity was normalized for movement speed effects using linear regression to assess the decay of the signal. We used a within-subject design, and comparisons between Test and Sample sessions were made after assessing normality through the Kolmogorov-Smirnov test and Q-Q plots; t-tests or Wilcoxon tests were applied accordingly. One-sample t-tests determined whether the DI or GLM coefficients significantly deviated from zero, with statistical significance set at p < 0.05.

## RESULTS

### Cholinergic activity is robustly correlated to the logarithm of the animal’s movement speed

First, we addressed whether cholinergic dynamics in the septo-hippocampal circuitry enable multiplexed coding of movement speed. Toward that goal, we measured the population activity of septo-hippocampal cholinergic projection neurons using fiber photometry in freely behaving mice performing a hippocampus-dependent ObLoM task (Figures 1A-C). Immunohistological data from coronal brain slices of the MSDB showed colocalization of GCaMP-positive neurons and ChAT-immunolabeled neurons, confirming that GCaMP expression was largely confined to cholinergic neurons in the MSDB (Example shown in Figures 1D-G).

**Figure 1.**
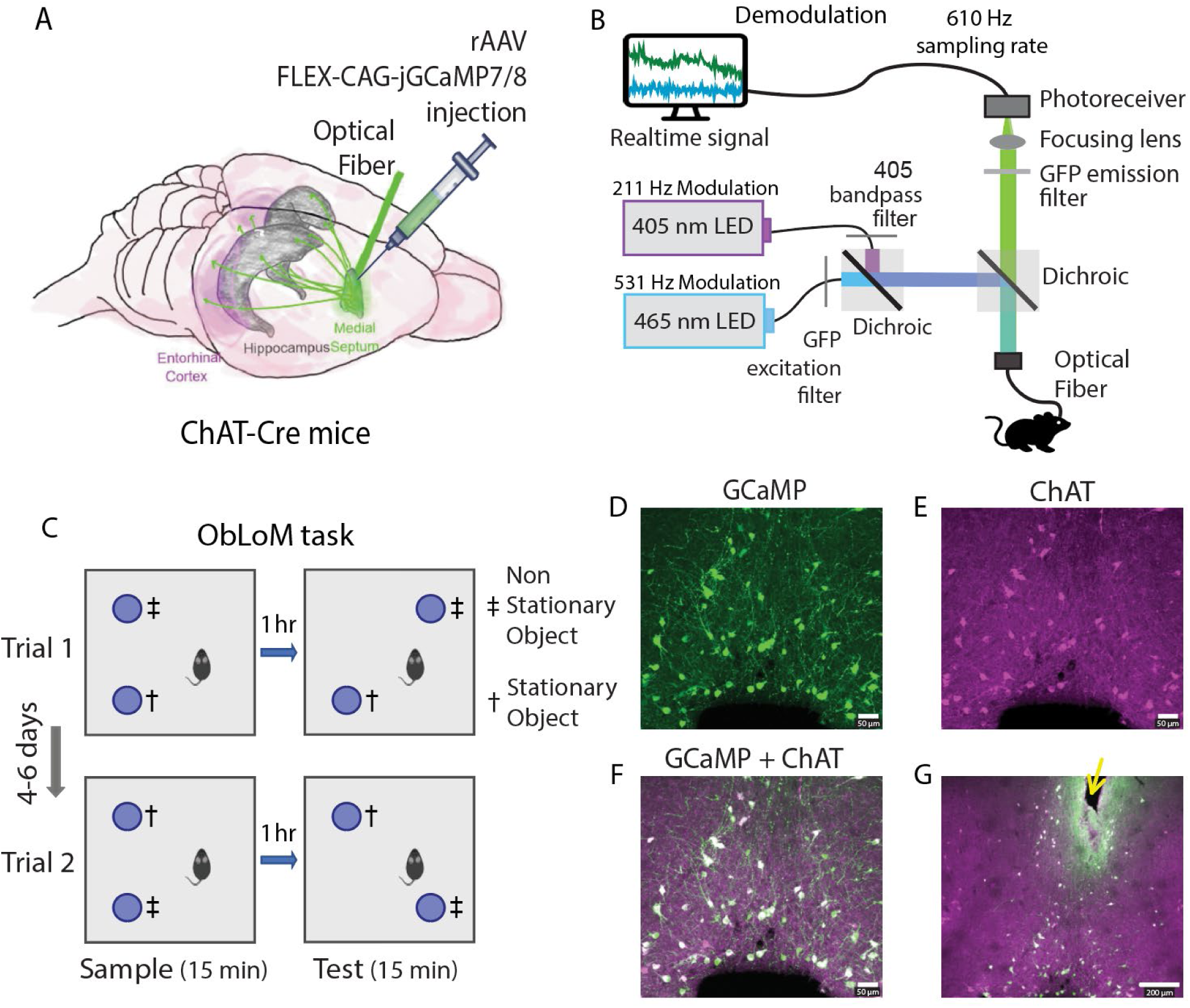
Experimental design. **A.** Schematic drawing of the surgical approach taken to perform fiber photometry of cholinergic septo-hippocampal projection neurons using jGCaMP7/8 in freely-behaving ChAT-Cre mice. **B.** Schematic drawing of the fiber photometry system. **C.** Design of the object location memory (ObLoM) task. During the Sample phase (15 minutes), mice explored two identical objects in a 40 x 40 cm² square arena with 30 cm high walls. During the Test phase (15 minutes), one of the two objects was moved to a novel location (Non-Stationary Object). The Sample and Test phases were separated by a 1-hour delay phase, during which the mouse was returned to its home cage (6 mice). Experiment was repeated as trial 2 in a counterbalanced design after 4-6 days (4 mice). **D-G.** Immunohistological verification of the optical fiber track position and cell type-specific expression of jGCaMP7/8 in cholinergic neurons of the MSDB. (D) Green color indicates jGCaMP7/8 fluorescence, (E) magenta color indicates immunolabeling of ChAT, a marker for cholinergic neurons. (F) White color indicates colocalization of jGCaMP7/8 fluorescence and ChAT immunostaining (Scale bar, 50 μm). (G) Histological confirmation of the fiber track position (yellow arrow points to tissue displaced by the implanted optical fiber) within the MSDB (Scale bar, 200 μm).

Previous experiments showed a strong correlation of cholinergic activity to the logarithm of the animal’s movement speed during free foraging (Kopsick et al., 2022). We therefore first asked whether the correlation of cholinergic activity to the logarithm of animal’s movement speed was altered by the engagement of the mice in the ObLoM task. As described previously (Kopsick et al., 2022), we computed the movement speed of the animal’s position using DeepLabCut (see Materials and Methods) and computed Pearson’s correlation coefficient. As during free foraging, cholinergic activity in mice performing the ObLoM task showed a strong linear correlation to the logarithm of the animal’s movement speed (Figures 2D-F), with striking similarity to the observed relationship of cholinergic activity to the animal’s movement speed found in freely ambulating mice (Kopsick et al., 2022). The correlation between cholinergic activity and the logarithm of movement speed was significantly higher than the correlation between cholinergic activity with the movement speed (R_Speed_ = 0.32 ± 0.15; R_Log2(speed)_ = 0.38 ± 0.18; mean ± Std; t=-7.72, p=<.001, Paired t-test) (Figures 2A, 2D).

**Figure 2.**
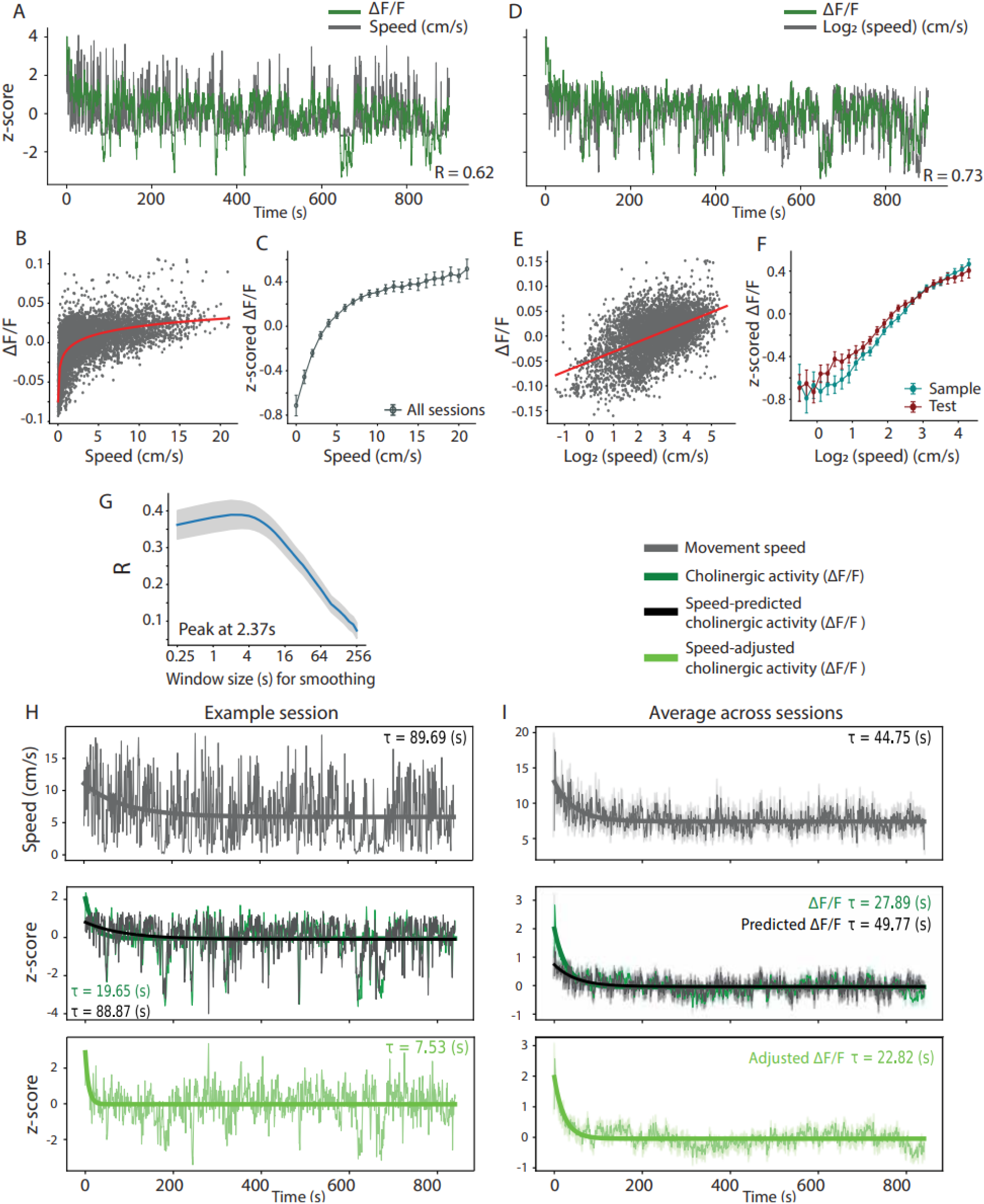
Cholinergic activity is linearly correlated to the logarithm of the animal’s movement speed during an object location-memory task. **A.** Correlation between cholinergic activity, quantified as ΔF/F, and the animal’s movement speed, for one example session recorded during the Test phase. Signals are smoothed with a 1-second window; R, Pearson’s correlation coefficient. **B.** Scatter plot of cholinergic activity, quantified as ΔF/F, as a function of the animal’s movement speed for an example session, with an exponential function fitted to the data (red). **C.** Z-scores of ΔF/F of cholinergic activity across all sessions as a function of the animal’s movement speed (20 sessions). Speed is binned with a bin width of 1 cm/s. Data shown as mean ± s.e.m. **D.** Same data as in (A) but using the logarithm of the animal’s movement speed. **E.** Same data as in (B) but using the logarithm of the animal’s movement speed, with a linear function fitted to the data (red). **F.** Z-scores of ΔF/F of cholinergic activity across Test sessions (red, 10 sessions) and Sample sessions (blue, 10 sessions) as a function of the animal’s movement speed. Speed is binned with a bin width of 0.2. Data shown as mean ± s.e.m. **G.** Mean ± s.e.m. of time scale-dependent Pearson correlation coefficient distributions between the smoothed cholinergic activity and the logarithm of the animal’s movement speed (20 sessions). **H-I.** Time-series data on movement speed and cholinergic activity, quantified as ΔF/F, from one example session (H) and averaged across all sessions (I). Time-series data are smoothed with a 1-second window for better illustration. Thick curves show the exponential fit to the data; τ = time constant of exponential fit. Data in (I) shown as mean ± s.e.m.; n = 20 sessions.

Furthermore, we employed a LMM to examine differences in the linear correlation between cholinergic activity and the logarithm of movement speed during the Sample and Test phases of the ObLoM task (Figure 2F). The analysis revealed a significantly lower slope during the Test sessions, though the effect size was small (speed × phase coefficient = −0.06; *z* = −30.19; *p* < 0.001; 95% CI = [−0.064, −0.056]).

Next, we asked what the optimal timescale is for cholinergic activity to encode movement speed. After applying smoothing window sizes ranging from 0.25 s to 256 s to the cholinergic signal (see Materials and Methods), we observed that the correlation between the logarithm of the animal’s movement speed and the cholinergic activity was maximal at a timescale of 2.37 s (Figure 2G), similar to what have been previously observed during free foraging behavior (Dannenberg et al., 2016). Taken together, these data suggest that the animal’s movement speed is robustly encoded by cholinergic activity in the septo-hippocampal system, despite the engagement of animals in a spatial memory task.

### Exponentially decaying cholinergic activity signals environmental novelty

In addition to encoding movement speed, cholinergic activity in the basal forebrain is critically involved in regulating arousal (Acquas et al., 1996; Dannenberg et al., 2016; Marrosu et al., 1995) and spatial novelty (Aloisi et al., 1997; Bianchi et al., 2003; Giovannini et al., 2001). However, whether novelty-triggered increases in cholinergic activity are tonic or phasic remains unknown. Here, we leveraged the sub-second temporal resolution of fiber photometry to investigate the rise and decay times of cholinergic activity when the animal is moved from its home cage to the experimental arena. During the transition, we observed an instantaneous increase in cholinergic activity that showed a slow decay of 28s (Figure 2I, middle plot). Since cholinergic activity increases with the animal’s movement speed, we asked whether the sharp increase followed by a slow decay in cholinergic activity does in fact signal environmental novelty or is simply due to increased movement speed at the beginning of the Sample and Test sessions. To answer this question, we fitted exponential decay functions to both the movement speed signal and the cholinergic activity and compared the decay constants (tau). While both movement speed and cholinergic activity were elevated during the initial minutes of each session, the decay time for movement speed was longer than the decay time for cholinergic activity (45-s vs. 28-s), suggesting that the change in cholinergic activity cannot be explained by changes in movement speed. To further corroborate that the elevated cholinergic activity at the beginning of the Sample and Test phases codes for environmental novelty, we adjusted the observed cholinergic activity for the effect of movement speed using a linear regression. The movement speed-adjusted cholinergic activity was substantially increased at the beginning of Sample and Test sessions, with an average decay time of 23 seconds. This data suggests that cholinergic activity encodes both movement speed and environmental novelty in parallel at different timescales.

### Mice learn and recall the location of objects in the object location memory task

To test whether cholinergic signals differentiate between the exploration of objects in novel vs. familiar locations, we recorded cholinergic dynamics during both the Sample and Test phases of the ObLoM task. We first assessed task performance using the DI (see Methods), analyzing data in 3-minute intervals for both the Sample and Test sessions (Figure 3A). The differences in DI values between the Test and Sample sessions were significantly different from zero only in the first 3-minute interval (p = 0.008; t = 3.4; One-Sample t-test). During this initial period, mice spent significantly more time exploring the object placed in the novel location than the object in the familiar location during the Test phase (Figure 3B). However, the timescales at which cholinergic signaling in the septo-hippocampal system is associated with behavioral events and the learning of novel object-space associations are unknown.

**Figure 3.**
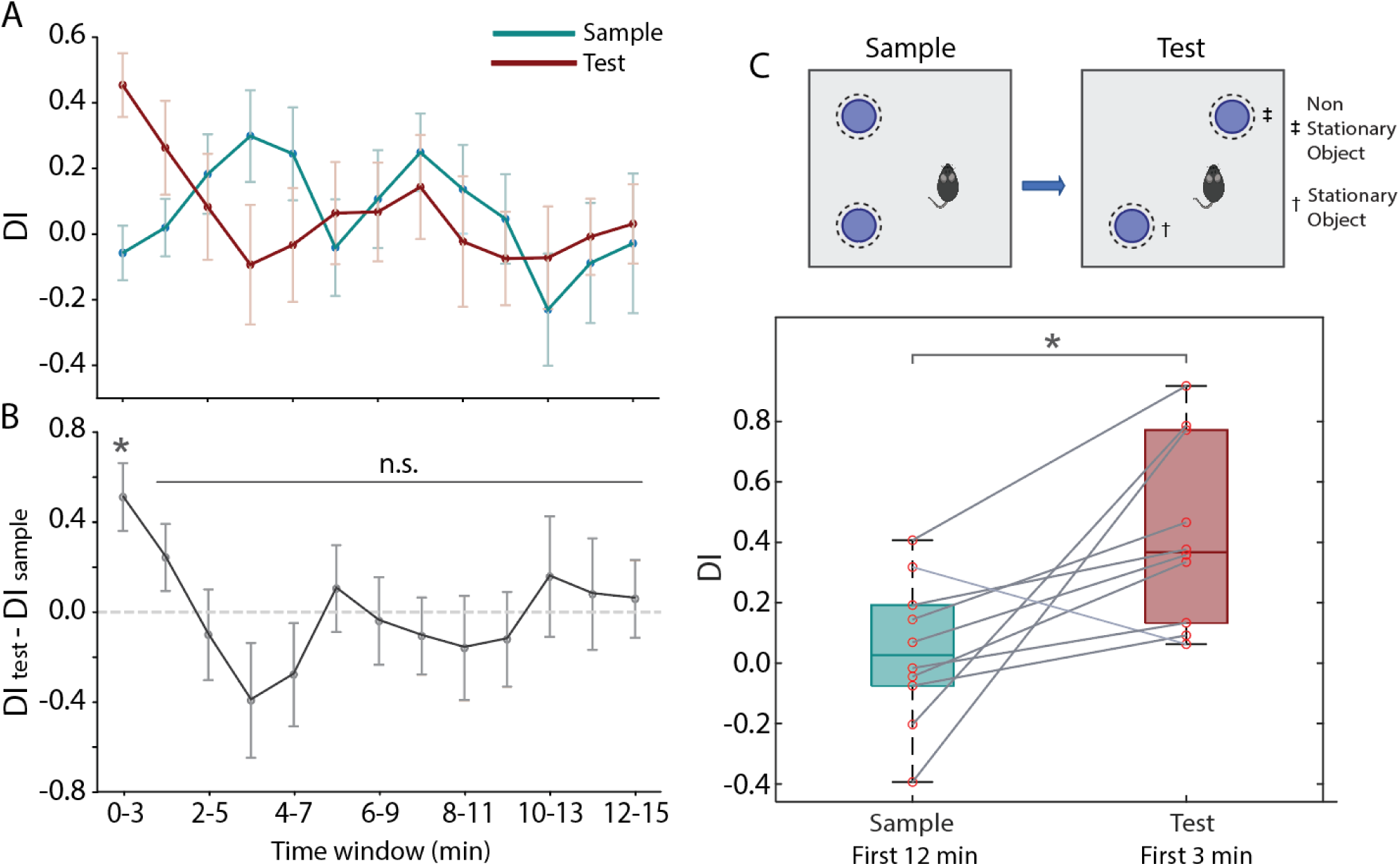
Mice recall the novel location of a non-stationary object in the ObLoM task. **A.** DI for Sample (blue) and Test (red) sessions computed for 3-minute intervals using a sliding window. Data show mean ± s.e.m. **B.** Difference in DI between Test and Sample sessions computed for 3-minute intervals using a sliding window; *p = 0.008; n.s. = not significant; n = 10 sessions. **C.** *(Top)* Illustration of the object location memory task (ObLoM). *(Bottom)* Data on DI for Sample and Test sessions, computed from data on the first 12-min and 3-min, respectively. *p = 0.016; n =10. DI = discrimination Index.

Based on these findings and consistent with the literature, we used data from the first 3 minutes of the Test session to compute the DI, while using data from the first 12 minutes of the Sample session to compute the DI under baseline conditions. This approach aligns with the methodology used in previous studies, facilitating comparisons across research studies (Schlesiger et al., 2021). During the Test but not the Sample session, mice consistently devoted more time to exploring the non-stationary object compared to the stationary object (DI_Sample_ = 0.04 ± 0.24; DI_Test_ 0.43 ± 0.3; mean ± Std), resulting in a significant difference in object discrimination between the Sample and Test phases (t = −2.97; p = 0.016; Paired t-test) (Figure 3C)

### Phasic cholinergic activity signals novelty of object locations and is correlated to behaviors associated with memory-guided navigation

We next investigated whether phasic activity in the cholinergic signal encodes the novelty of an object location. Because cholinergic activity is widely recognized as an important modulator of neural activity underpinning memory-guided navigation, we quantified cholinergic signals associated with exploring objects at familiar and novel locations in relation to cholinergic signals associated with the logarithm of movement speed—correlated with locomotion—as well as grooming and rearing. Because of the nature of these behaviors, only object exploration could overlap in time with rearing or locomotion (Figure 4A-B). To compare the effects of each behavior on cholinergic activity, we applied a GLM to the entire 15-minute duration of each session (10 Sample sessions, 10 Test sessions) and quantified the impact of behavioral states during both Sample and Test sessions (see Table 3) (Figure 4C). The results showed that the logarithm of movement speed exhibited a robust positive effect on cholinergic activity in both Sample and Test sessions; rearing did not significantly impact cholinergic activity in either Sample or Test sessions; and grooming exhibited a negative effect on cholinergic activity in both Sample and Test sessions.

**Figure 4.**
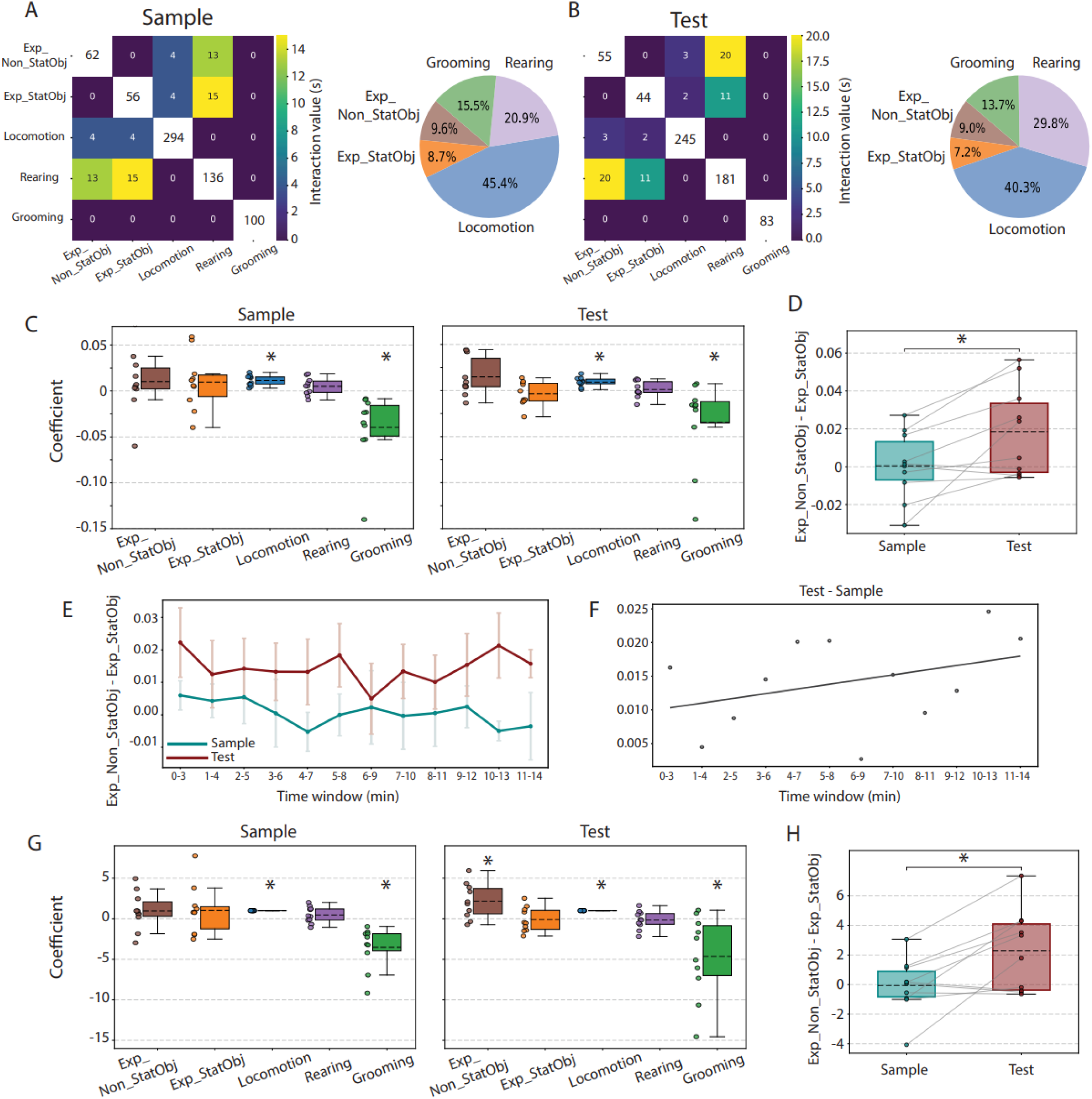
Phasic cholinergic activity signals novelty of object locations and are correlated to behavioral states. **A.** *(Right)* Average distribution and co-occurrence of time spent by animals in different behavioral states during Sample sessions (10 sessions). *(Left)* Pie charts display the percentage of each behavior relative to the others. **B.** Same as (a) but for Test sessions (10 sessions). **C.** Summary data comparing the effects of different behavioral states on cholinergic activity during Sample (left, 10 sessions) and Test (right, 10 sessions) sessions. Coefficients are extracted from general linear model (GLM) results; * Significantly different from zero. **D.** Differences in the effects of exploring non-stationary versus stationary objects on cholinergic activity during Test (red) and Sample (blue) sessions. Coefficients are extracted from GLM results; * Significantly different. **E.** Mean ± s.e.m. of the differences in the effect of object exploration on cholinergic activity, calculated over 3-minute intervals using a sliding window for Sample (blue, 10 sessions) and Test (red, 10 sessions) sessions. Coefficients are extracted from the applied GLM over 3-minute intervals, including only periods with at least 1 second of object exploration. **F.** Linear regression analysis of the differences in data shown in (E) between Test and Sample sessions. **G.** Same data as in (B) after normalizing by the effect of movement speed. **H.** Same data as in (E) after normalizing by the effect of movement speed.

**TABLE 3.** Results of One-Sample t-test (Test Value = 0) for each coefficient in sample versus Test sessions, before and after normalizing to speed coefficients.

| Sample sessions |  |  |  |  | Test sessions |  |  |  |
| --- | --- | --- | --- | --- | --- | --- | --- | --- |
| Coefficient | Mean ± Std | t | P-value | 95% CI | Mean ± Std | t | P-value | 95% CI |
| Exp_Non_StatObj | 0.0102 ± 0.034 | 0.944 | 0.37 | [-0.0142, 0.0345] | 0.0149 ± 0.021 | 2.25 | 0.051 | [0, 0.03] |
| Exp_StatObj | 0.0097 ± 0.0307 | 0.995 | 0.345 | [-0.0123, 0.0317] | -0.0035 ± 0.0133 | -0.823 | 0.432 | [-0.013, 0.006] |
| Log<sub>2</sub>(Speed) | 0.0114 ± 0.0057 | 6.3 | <.001 | [0.01, 0.02] | 0.0091 ± 0.0051 | 5.67 | <.001 | [0.01, 0.01] |
| Rearing | 0.005 ± 0.0095 | 1.658 | 0.132 | [-0.0018, 0.0118] | 0.0012 ± 0.0089 | 0.4184 | 0.685 | [-0.0052, 0.0076] |
| Grooming | -0.0397 ± 0.0388 | -3.23 | 0.01 | [-0.07, -0.01] | -0.0344 ± 0.0475 | -2.29 | 0.048 | [-0.07, 0] |
| **After normalizing to speed** |  |  |  |  |  |  |  |  |
| Exp_Non_StatObj | 0.9626 ± 2.3526 | 1.293 | 0.228 | [-0.7202, 2.6454] | 2.1845 ± 2.1674 | 3.1873 | 0.011 | [0.6342, 3.7349] |
| Exp_StatObj | 1.0306 ± 3.0497 | 1.068 | 0.313 | [-1.1509, 3.2121] | -0.1007 ± 1.5164 | -0.21 | 0.838 | [-1.1854, 0.984] |
| Log<sub>2</sub>(Speed) | 1 ± 0 | Infinity | <.001 | [1, 1] | 1 ± 0 | Infinity | <.001 | [1, 1] |
| Rearing | 0.4537 ± 1.0021 | 1.431 | 0.186 | [-0.2631, 1.1704] | -0.1553 ± 1.1413 | -0.4302 | 0.677 | [-0.9717, 0.6611] |
| Grooming | -3.5262 ± 2.6191 | -4.257 | 0.002 | [-5.3997, -1.6527] | -4.6617 ± 5.1056 | -2.8874 | 0.018 | [-8.3137, -1.0096] |

We next asked whether phasic cholinergic activity encodes the novelty of an object-space association. During the Sample session, both the non-stationary and stationary object are novel to the animal. In contrast, during the Test session, the non-stationary object is now found at a novel location, while the stationary object is still found at its familiar location. To test whether cholinergic activity reflects the encoding of the novel object location, we computed the effect of exploring stationary versus non-stationary object on cholinergic activity in both Test and Sample phases. We then compared Sample and Test sessions (Figure 4D). The difference during Test sessions was significantly higher than in Sample sessions (Test: 0.0005 ± 0.0177; Sample: 0.0184 ± 0.0239; mean ± Std; t = −3.05; p = 0.014; Paired t-test), suggesting that phasic cholinergic activity supports the encoding of a novel object-place association.

Our analysis of behavioral data showed that, while mice initially spent more time exploring the object at the novel location, they soon adopted a pattern of exploring both object locations equally (Figures 3A-B). We asked whether this initial object preference in exploration behavior was mirrored by cholinergic dynamics (Figures 4E-F). We applied GLM over 3-minute intervals using a sliding window across sessions. Our results showed that cholinergic activity continued to differentiate between objects at novel and familiar locations throughout most of the 15-minute recording session (y-intercept = 0.0096; t = 2.367; *p* = 0.039; 95% CI = [0.001, 0.019]; slope = 0.0007; t = 1.265; *p* = 0.235; 95% CI = [−0.001, 0.002]; not significantly different than zero). These findings suggest that phasic cholinergic signals convey a spatial novelty signal even when differences in behavioral patterns are no longer evident.

In fiber photometry, variability across recording sessions arises from several technical factors, such as fluctuations in laser power, inconsistent fiber coupling, and changes in signal strength with changing expression levels of the fluorophore (Simpson et al., 2024). These factors can influence both baseline fluorescence and signal amplitude, and hence the signal-to-noise ratio, making direct session-to-session comparisons difficult. To improve the interpretability of data averaged across sessions, we implemented a normalization approach that adjusted model coefficients to a common reference. Specifically, we used the coefficient for movement speed as a reference because of the robust and strong correlation of movement speed to cholinergic activity across all mice and sessions (Figure 4G). Normalizing the model parameters to the effect of movement speed provided a robust framework for comparing signals across recording sessions and experimental conditions. The results confirmed our previous conclusions (see Table 3). Concretely, rearing did not significantly impact cholinergic activity in either Sample or Test sessions, and grooming exhibited a negative effect on cholinergic activity. The difference in the effect of exploring objects in familiar versus novel locations on cholinergic activity during Test sessions was significantly higher than in Sample sessions (Test: 2.2852 ± 2.7216; Sample: −0.068 ± 1.8568; mean ± Std; t = −3.11; p = 0.012; Paired t-test) (Figure 4H), confirming our previous conclusion that phasic cholinergic activity encodes the novelty of an object location.

### Changes in cholinergic activity before, during, and after behaviors associated with memory-guided navigation

Having established that phasic cholinergic activity codes for various behaviors and provides a novelty signal for object locations, we next quantified the event-triggered temporal profile of cholinergic activity, including the moments before and after the onset and offset of a given behavior. Toward that end, we aligned cholinergic signals with behavioral transitions into and out of each behavior, including exploration of objects, covering a window from 5 seconds before the onset to 5 seconds after the offset of a given behavior (see Materials and Methods). Since the duration of individual behavioral bouts differed, we linearly time-warped behavioral bout durations between the onset and offset of bouts, so that cholinergic activity could be averaged across behavioral bouts, revealing the temporal profile of cholinergic dynamics associated with each behavior (total of 10 Sample and 10 Test sessions from 6 mice). Then, we used movement speed to predict cholinergic activity and compared it to the observed cholinergic signal.

Interestingly, cholinergic activity transiently increased prior to the onset of locomotion and decreased after its termination. The timing of the increase in the observed cholinergic activity (reaching 50%) was 0.47 seconds earlier than the predicted cholinergic signal based on movement speed, potentially reflecting a preparedness to move. Furthermore, as shown previously (Zhang et al., 2021), cholinergic activity displayed a decrease during grooming, where this decrease was synchronous with the predicted cholinergic signal derived from movement speed. For rearing, cholinergic activity remained relatively constant and was notably higher than the predicted cholinergic signal, suggesting that cholinergic activity during rearing may be influenced by factors beyond movement signals, potentially reflecting a distinct cognitive or exploratory component (Figure 5A).

**Figure 5.**
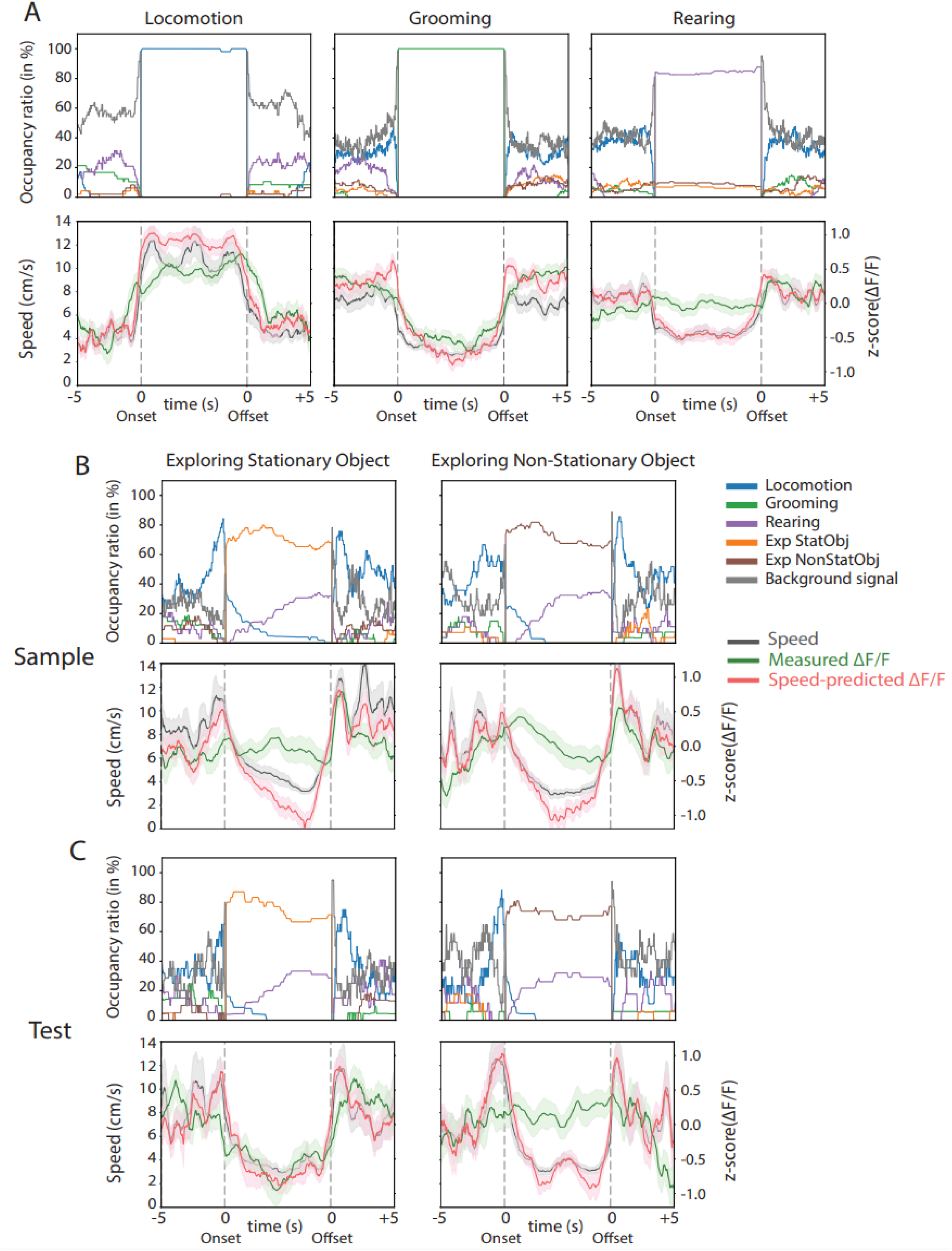
Fast cholinergic transients across cognitive and behavioral states. **A.** *(Top Row)* Occupancy ratios of behavioral states 5 s before the onset, during, and 5 s after the offset of locomotion, grooming, and rearing events. Only events with no occurrence of the behavioral state in question during the 4 s before or after the onset and offset were analyzed. Between the onset and offset of a behavioral state, data are plotted on a relative timescale. Different colors represent distinct behaviors: locomotion (blue), grooming (green), rearing (purple), exploratory behaviors associated with stationary objects (orange), exploratory behaviors associated with non-stationary objects (brown), and background (gray) indicating the absence of all other behaviors. *(Bottom Row)* Data on movement speed (black), observed cholinergic activity (green), and cholinergic activity predicted from movement speed (red). Data show mean ± s.e.m. Locomotion events, n = 47; Grooming events, n = 73; Rearing events, n = 85. Data from 6 mice. **B.** Data on exploring the stationary and non-stationary objects in the *Sample* session. Data are visualized in the same way as in (A). Exploring the stationary object, n = 32; exploring the non-stationary object, n = 27. **C.** Data on exploring the stationary and non-stationary objects in the *Test* session. Data are visualized in the same way as in (B). Exploring the stationary object, n = 20; exploring the non-stationary object, n = 17.

With respect to the function of cholinergic modulation in the context of providing a novelty signal for object locations, the temporal profiles show that cholinergic activity remains high during the exploration of objects in Sample sessions, where both objects are novel, as well as during the exploration of non-stationary object at novel location in Test sessions. In all of these cases, the observed cholinergic signal is higher than the predicted signal based on movement speed. However, cholinergic activity noticeably decreased while exploring the stationary object at familiar location during Test sessions, almost synchronously with the predicted cholinergic signal (Figure 5B).

## DISCUSSION

The primary objective of this research was to quantify the extent and timescales at which different cognitive and behavioral states correlate with the activity of medial septal cholinergic neurons in the context of a spatial memory task. The results demonstrate that cholinergic activity encodes movement speed, the novelty of object locations, and environmental novelty in parallel but at different timescales. Furthermore, we show fast (sub-second to seconds) cholinergic transients correlate with the initiation and termination of different behaviors.

Previous studies of cholinergic signals in cortical regions have shown that cholinergic modulation can operate at different timescales, with cortical release occurring in either tonic or phasic modes (Disney and Higley, 2020; Sarter and Lustig, 2020). Concretely, phasic signals play a critical role in processing temporally discrete events, such as sensory stimulation and cue detection (Gritton et al., 2016; Zhu et al., 2023) and sensorimotor integration (Yogesh and Keller, 2024; Zou et al., 2024). Similarly, phasic cholinergic signaling has been implicated in the coding of movement speed both during locomotion and in stationary mice (Dannenberg et al., 2016; Kopsick et al., 2022). Phasic signals may be facilitated by ACh release at synaptic terminals, as opposed to volume transmission, supported by anatomical studies using electron microscopy and super-resolution imaging (Muller et al., 2013; Takács et al., 2018; Turrini et al., 2001) showing cholinergic axons form synaptic contacts. In contrast, tonic signals have been shown to support cognitive processes, such as learning and memory (Dannenberg et al., 2017; Hasselmo, 2006). In particular, high levels of cholinergic tone are causally linked to the “online” theta state in awake, exploratory mice (Dannenberg et al., 2015; Vandecasteele et al., 2014) and during rapid eye movement (REM) sleep (Marrosu et al., 1995), and facilitate synaptic plasticity and hippocampal encoding of spatial and contextual information (Záborszky et al., 2018). Conversely, low cholinergic tone during slow-wave sleep and SPW-Rs provides a permissive environment for the replay of neuronal activity patterns (Zhang et al., 2021). However, whether septo-hippocampal cholinergic modulation uses multiplexing of fast and slow signals to support the encoding of novel spatial information, movement speed, and behaviors has remained elusive. Here, we observed a strong correlation between cholinergic activity and the brief moments during which a mouse explores a novel object location while also coding for the logarithm of movement speed. These correlations remained high at a sub-second timescale, indicating a fast or phasic mode of cholinergic signaling. At the same time, we observed a rapid increase but slow decay in cholinergic activity at each start of a session when animals transitioned from the home cage to the test arena—a new environment. Using a linear regression model to adjust for the effects of movement speed on cholinergic activity, we quantified the decay time constant as ∼23 seconds, supporting the hypothesis that ACh release in the hippocampus signals environmental novelty in addition to coding for movement speed and novel object locations.

Taken together, the results on multiplexed fast and slow cholinergic dynamics associated with exploration of novel object locations and novel environments support the theory that ACh release in the hippocampus and related areas supports the encoding of spatial memories at both short and long timescales (Hasselmo, 2006; Hasselmo et al., 1995; Zhang et al., 2021). Moreover, they are consistent with previous findings on the role of cholinergic activity in cognitive processes such as arousal (Acquas et al., 1996), novelty detection (Acquas et al., 1996), memory encoding (Hasselmo, 2006), attention (Hasselmo and McGaughy, 2004; Peters et al., 2011), and sensory integration (Kuchibhotla et al., 2017). In particular, coordination of cholinergic activity between hippocampal and neocortical regions may enhance sensory input processing to facilitate the encoding of novel spatial information.

During the first three minutes of the Test but not Sample sessions, mice spent significantly more time exploring the non-stationary object at the novel location than the stationary one. This behavior demonstrates that mice have learned and recalled object locations in the object location memory task. A previous study demonstrated that cholinergic signals are strongly correlated with body movements while the mouse is stationary (Kopsick et al., 2022). We therefore employed a GLM to distinguish the effect of movement speed, rearing, grooming, and exploring objects on cholinergic activity; we observed a significantly stronger effect of exploring object at novel location on the cholinergic signal compared to exploring object at familiar location. Intriguingly, the phasic cholinergic signals conveying the spatial novelty signal were present throughout the entire session, outlasting the first three minutes where differences in behavioral exploration patterns were evident. These data suggest that phasic cholinergic signals support the learning of novel object-place associations rather than driving novelty-related behavior.

While the effects of arousal and novelty on behavior and information processing are closely linked, there is a subtle yet important distinction. Novelty refers to the unfamiliarity or “newness” of a stimulus which triggers an orienting response facilitating sensory integration, memory encoding, and learning (Barto et al., 2013; Tanner et al., 2024; Weierich et al., 2010). Arousal, on the other hand, represents a broader state of heightened neural and physiological activity that prioritizes attention and sensory integration for relevant stimuli and applies to various affective stimuli, regardless of their novelty (Weierich et al., 2010). Although novelty often leads to arousal, arousal can also arise from factors unrelated to novelty, such as emotional valence or sensory intensity (Barto et al., 2013; Tanner et al., 2024; Weierich et al., 2010). Future studies are needed to tease apart the differential contributions of arousal and spatial novelty to cholinergic signaling in the hippocampal formation to help elucidate the mechanisms by which cholinergic signaling supports the integration of sensory and behavioral information during spatial memory tasks.

Cholinergic activity is widely recognized as an important modulator of neural activity underpinning memory-guided navigation (Dannenberg et al., 2016; Hinman et al., 2018). We therefore investigated the temporal dynamics of cholinergic activity preceding, during, and succeeding prominent behaviors typically associated with the encoding, retrieval, and consolidation of spatial information such as locomotion, rearing, and grooming. These analyses revealed that cholinergic activity was significantly reduced during grooming in both Sample and Test sessions, consistent with data from a previous study on decreased ACh release in the hippocampus during SPW-Rs, which predominantly occur during grooming periods in the awake animal (Zhang et al., 2021). In contrast, rearing showed no statistically significant impact on cholinergic activity in the context of the ObLoM task. This negative finding suggests that cholinergic activity may not be correlated to rearing events in a highly familiar environment. However, when analyzing the temporal profile of cholinergic activity during rearing events, we found cholinergic activity was higher than predicted by the low movement speed, consistent with findings of increased cholinergic activity during rearing episodes in the context of free exploration (Kopsick et al., 2022) and with results from a different lab demonstrating increased rearing duration but no changes in rearing frequency when optogenetically inhibiting medial septal cholinergic neurons of rats during an 8-arm win-shift paradigm (Cassity et al., 2023). The latter findings were attributed by the authors to animals needing extra time to encode novel information when acetylcholine release is reduced. Investigating the requirement of cholinergic signals for integrating sensory information during rearing events would be a worthwhile endeavor for future studies.

To our surprise, we found an increase in cholinergic activity *before* the onset of locomotion. The rise in cholinergic activity before the onset of motor actions suggests that cholinergic signals prepare neural circuits in the hippocampus for the processing of movement-related information, potentially facilitating sensorimotor integration during spatial navigation or exploratory behavior. Such a function of cholinergic modulation within the septo-hippocampal circuitry aligns well with cholinergic mechanisms observed in other cortical regions, such as the necessity of Ach in the auditory cortex for signaling behavioral context transitions (Kuchibhotla et al., 2017) and the early activation of nicotinic acetylcholine receptors (nAChRs) in the prefrontal cortex, which is crucial for rapid orienting and cue detection (Howe et al., 2017). The increase in cholinergic activity before the onset of locomotion is consistent with the hypothesis that cholinergic modulation promotes a brain state optimized for the integration of sensory information, such as sensory flow during movement, and self-motion codes during locomotion. This idea is consistent with findings that faster and longer treadmill running elicits larger ACh responses in visual cortex (Neyhart et al., 2024). Future research involving continuous cholinergic signal recordings during object location memory tasks in both light and darkness is needed to clarify the role of optic flow in cholinergic modulation. Additionally, optogenetic approaches targeting cholinergic neurons during various behavioral states may help quantify to what extent cholinergic modulation is required for sensory integration and motor tasks.

In the context of novel object exploration, cholinergic activity showed distinct patterns. It remained high during the exploration of objects in Sample sessions and the non-stationary object at a novel location in Test sessions, despite reduced movement speed. In contrast, it decreased during the exploration of the stationary object at a familiar location in Test sessions, closely aligning with movement speed dynamics. The difference in the temporal profiles of cholinergic activity between an animal exploring a novel versus a familiar object location during the Test phase cannot be attributed to any potential object, place, or object-place preference because i) the objects were identical, ii) the temporal profiles in cholinergic activity associated with exploring the stationary and non-stationary objects did not differ in the Sample session, and iii) we used a counter-balanced task design with respect to which of the two objects were moved to a novel location. Moreover, the magnitudes and temporal profiles of phasic cholinergic activities associated with exploration of both stationary and non-stationary objects in the Sample phase are similar to the magnitude and temporal profile of cholinergic activity associated with exploration of the object moved to a novel location in the Test phase. Collectively, these data are consistent with ACh providing a novelty signal for spatial location, potentially promoting the learning of novel object-space associations.

To summarize, our results demonstrate multiplexing of cholinergic signals in the septo-hippocampal system across multiple timescales. Concretely, slow changes in cholinergic activity correlate with environmental novelty, and phasic cholinergic activity correlates with the exploration of novel object locations, as well as movement speed and changes in cognitive and behavioral states. These data suggest that cholinergic modulation of hippocampal circuits provides novelty signals facilitating the acquisition of spatial memories and enables the integration of sensory information and self-motion codes during spatial navigation. Multiplexed coding in the septo-hippocampal system aligns with mechanisms of cholinergic modulation observed in other cortical regions such as the prefrontal cortex, where ACh facilitates cue detection via fast-acting nicotinic ACh receptors and decision-making via slow-acting muscarinic ACh receptors (Howe et al., 2017), as well as the auditory cortex, where ACh supports sensory integration and context-dependent behavior (Kuchibhotla et al., 2017). Whether, and if yes, to what extent, fast and slow cholinergic dynamics in the hippocampal system mediate their effects on behavior and neural dynamics via differentially activating nicotinic and muscarinic ACh receptors, remains to be studied.

Overall, this study deepens our understanding of the mechanisms by which cholinergic dynamics in hippocampal memory circuits contribute to the coding of spatial novelty and spatial memories as well as the integration of sensory information and self-motion codes. The presented data will enable the evaluation of long-standing hypotheses regarding the function of cholinergic modulation in memory and exploratory behavior.

## Acknowledgements

This work was supported by the National Institute of Neurological Disorders and Stroke and the National Institute on Aging of the National Institutes of Health, grant numbers R00NS116129 and R21AG087912. The authors have no conflicts of interest.

